# Alpha-1 antitrypsin inhibits SARS-CoV-2 infection

**DOI:** 10.1101/2020.07.02.183764

**Authors:** Lukas Wettstein, Carina Conzelmann, Janis A. Müller, Tatjana Weil, Rüdiger Groß, Maximilian Hirschenberger, Alina Seidel, Susanne Klute, Fabian Zech, Caterina Prelli Bozzo, Nico Preising, Giorgio Fois, Robin Lochbaum, Philip Knaff, Volker Mailänder, Ludger Ständker, Dietmar Rudolf Thal, Christian Schumann, Steffen Stenger, Alexander Kleger, Günter Lochnit, Konstantin Sparrer, Frank Kirchhoff, Manfred Frick, Jan Münch

## Abstract

Severe acute respiratory syndrome coronavirus 2 (SARS-CoV-2) causes coronavirus disease 2019 (COVID-19). To identify factors of the respiratory tract that suppress SARS-CoV-2, we screened a peptide/protein library derived from bronchoalveolar lavage, and identified α1-antitrypsin (α1-AT) as specific inhibitor of SARS-CoV-2. α1-AT targets the viral spike protein and blocks SARS-CoV-2 infection of human airway epithelium at physiological concentrations. Our findings show that endogenous α1-AT restricts SARS-CoV-2 and repurposes α1-AT-based drugs for COVID-19 therapy.

## Main

SARS-CoV-2 is mainly transmitted through inhalation of contaminated droplets and aerosols and consequently infects cells of the respiratory tract. In most cases, infection is limited to the upper airways resulting in no or only mild symptoms. Severe disease is caused by viral dissemination to the lungs ultimately resulting in acute respiratory distress syndrome, cytokine storm, multi-organ failure, septic shock, and death. The airway epithelium acts as a frontline defense against respiratory pathogens, not only as a physical barrier and through the mucociliary apparatus but also via its immunological functions^1^. The epithelial lining fluid is rich in innate immunity peptides and proteins with antibacterial and antiviral activity, such as lysozyme, lactoferrin or defensins^2^. Currently, our knowledge about innate immune defense mechanisms against SARS-CoV-2 in the respiratory tract is very limited.

To identify endogenous antiviral peptides and proteins, we previously generated peptide/protein libraries from body fluids and tissues and screened the resulting fractions for antiviral factors^3^. This approach allowed to identify novel modulators of HIV-1^4,5^, CMV^6^ and HSV-2^7^ infection, with prospects for clinical development as antiviral drugs^8^. Here, we set out to identify factors of the respiratory tract that inhibit SARS-CoV-2. For this, we extracted polypeptides from 6.5 kg of homogenized human lung or 20 liters of pooled bronchoalveolar lavage (BAL), and separated them by chromatographic means. The corresponding fractions were added to Caco2 cells and infected with pseudoparticles carrying the SARS-CoV-2 spike protein^9^. None of the fractions of the lung library suppressed infection (Extended Data Fig. 1a). In contrast, fractions 42-45 of the BAL library prevented SARS-CoV-2 pseudoparticle infection almost entirely (Fig. 1a). The antiviral effect was comparable to that of 10 μM EK1, a spike-specific peptide fusion inhibitor used as control^10^ (Fig. 1a). Titration of BAL fractions 42 to 45 onto Caco2 cells confirmed that they prevent infection by SARS-CoV-2 spike pseudoparticles in a dose-dependent manner (Extended Data Fig. 2).

**Fig. 1.**
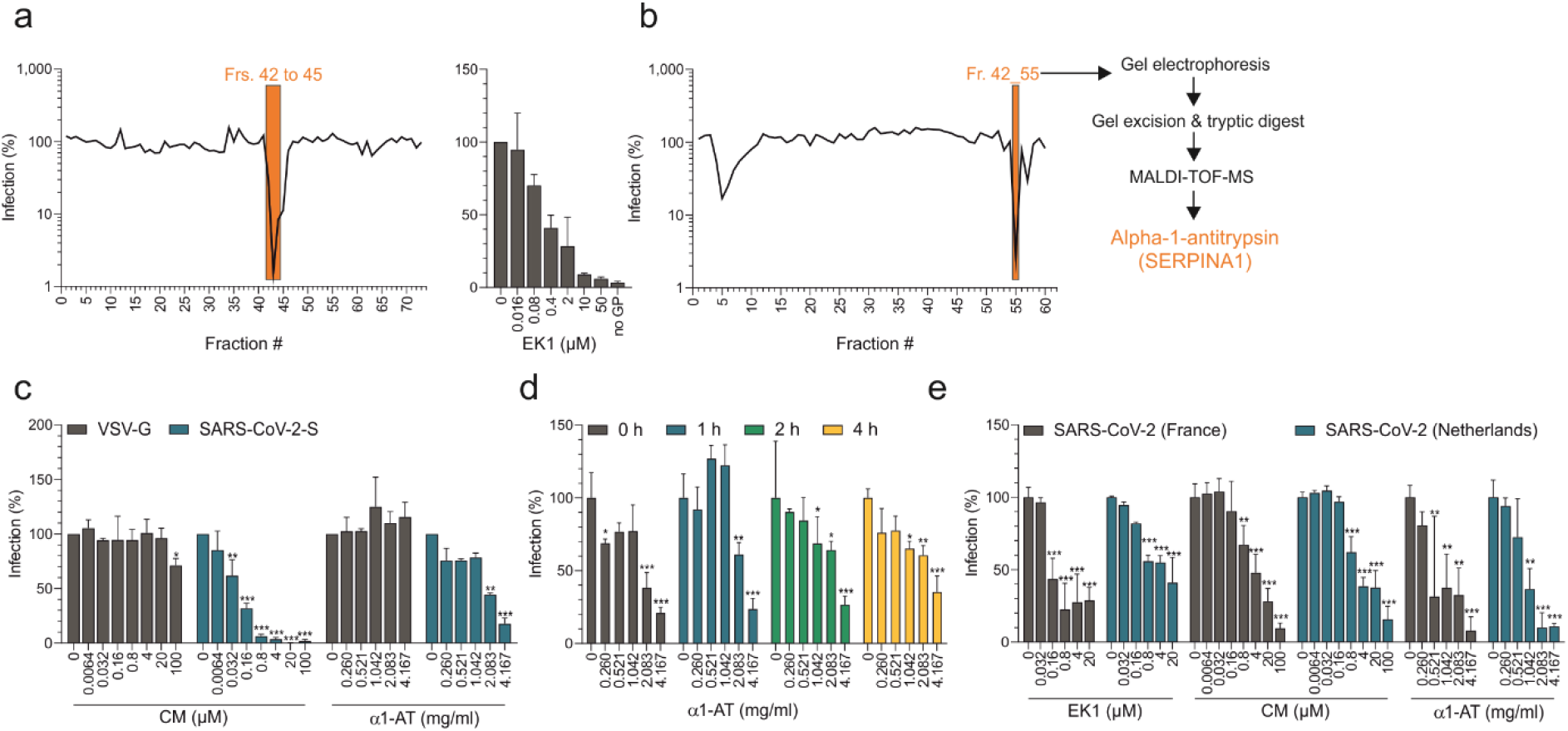
Identification of α1-AT as SARS-CoV-2 inhibitor. a) Fractions 42-45 of the bronchoalveolar lavage library and EK1 prevent SARS-CoV-2 spike pseudoparticle infection of Caco2 cells. b) Fraction 55 of further fractionated mother fraction 42 prevents SARS-CoV-2 spike pseudoparticle infection. Subsequent analysis of the antiviral fraction identified α1-AT as major component. c) α1-AT and the control small molecule inhibitor Camostate mesylate (CM) inhibit infection of pseudoparticles carrying SARS-CoV-2 spike but not VSV-G. d) α1-AT inhibits SARS-CoV-2 spike pseudoparticle infection at an early stage. CaCo2 cells were treated for indicated hours with α1-AT and then infected with SARS-CoV-2 spike pseudoparticles. e) α1-AT and controls block SARS-CoV-2 infection of TMPRSS2-expressing Vero E6 cells. Values shown in a and b are means from duplicate infections, values in c and d are means from two independent experiments performed in triplicates ± SEM, and values shown in e are means from triplicate infections ± SD.

To isolate the antiviral factor, the corresponding mother fraction 42 was further separated chromatographically and the resulting sub-fractions analyzed for antiviral activity. As shown in Fig. 1b, sub-fractions 42_4 to 42_9 and 42_57 reduced, and sub-fraction 42_55 almost completely prevented SARS-CoV-2 spike pseudoparticle infection. Analysis of these inhibitory fractions by gel electrophoresis showed distinct protein bands in sub-fractions 42_55 and 42_57, whereas no peptide/protein was detectable in sub-fractions 42_5 to 42_7 (Extended Data Fig. 3). The most active sub-fraction 42_55 contained a prominent band at ~ 52 kDa, which was also detectable in other active fractions but hardly present in those showing no antiviral activity (e.g. 42_49) (Extended Data Fig. 3). This band was excised from the gel and subjected to MALDI-TOF-MS revealing a 100 % sequence identity to α1-antitrypsin (SERPINA1) (Extended Data Fig. 4 and extended Table 1), a 52 kDa protease inhibitor^11^. The presence of α1-antitrypsin (α1-AT) in inhibitory fractions 42_55 and 42_57 was confirmed by Western Blots with an α1-AT-specific antibody (Extended Data Fig. 5).

To test whether α1-AT inhibits SARS-CoV-2 infection, we used a commercially available α1-AT preparation from human blood (Prolastin®). Camostat mesylate (CM), a small molecule inhibitor of the spike priming protease TMPRSS2 was included as control^9,12^. α1-AT and CM both suppressed SARS-CoV-2 spike pseudoparticle infection of Caco2 cells, with half-maximal inhibitory concentrations (IC_50_) of ~ 2 mg/ml for α1-AT, and ~ 0.05 μM for CM, respectively (Fig. 1c). Cell viability assays showed that α1-AT displays no cytotoxic effects at concentrations of up to 8.2 mg/ml, whereas CM reduced cell viability at concentrations of 200 μM, due to DMSO in the stock (Extended Data Fig. 6). The antiviral activity of α1-AT and CM was specific for the corona virus spike because infection of pseudoparticles carrying the G-protein of VSV was not affected (Fig. 1c). These data suggest that α1-AT may inhibit an early step in the SARS-CoV-2 replication cycle. In fact, time of addition experiments showed that α1-AT blocks infection if added to cells 1 to 4 hours prior to or during SARS-CoV-2 pseudovirus infection (Fig. 1d), but not if added 2 hours post infection (Extended Data Fig. 7).

To determine whether α1-AT inhibits not only Spike-containing pseudoparticles but also wild-type SARS-CoV-2, we examined its activity against two SARS-CoV-2 isolates from France and the Netherlands. For this, we assessed survival rates of Vero cells infected in the absence or presence of EK1, CM or α1-AT. In the absence of drugs, infection by both SARS-CoV-2 isolates resulted in a massive virus-induced cytopathic effect (CPE) and reduced cell viability by about 80% (Extended Data Fig. 8). Microscopic evaluation revealed the absence of CPE in the presence of high concentrations of the compounds (not shown). MTS assays confirmed a concentration-dependent inhibition of cell killing and viral replication by EK1 and CM (Extended Data Fig. 8, Fig. 1e) with average IC_50_ values against both SARS-CoV-2 isolates of 2.81 ± 2.52 μM for EK1, and 3.62 ± 0.03 μM for CM, respectively (Fig. 1e). Strikingly, α1-AT suppressed the French SARS-CoV-2 isolate with an IC_50_ of 0.58 mg/ml, and the Dutch strain with an IC_50_ of 0.81 mg/ml (Fig. 1e). Almost complete rescue of cell viability was observed at α1-AT concentration of 2-4 mg/ml (Extended Data Fig. 8, Fig. 1e).

To corroborate the anti-SARS-CoV-2 activity of α1-AT under conditions reflecting the in vivo conditions, fully differentiated human airway epithelial cells (HAEC) grown at air-liquid conditions were treated with 0.5 mg/ml of α1-AT and then infected with SARS-CoV-2. As control we used 5 μM of remdesivir, which has previously been shown to suppress coronavirus replication in HAECs^13^. At day 2, cells were fixed and stained with antibodies against SARS-CoV-2 spike^14^, and α-tubulin as marker for ciliated cells at the apical surface^15^. In infected mock-treated HAECs, SARS-CoV-2 spike expression was readily detectable, mostly in neighboring ciliated cells (Fig. 2), demonstrating that the epithelia were productively infected. Spike expression levels in α1-AT and remdesivir treated cultures were greatly reduced (Fig. 2). Thus, α1-AT suppresses SARS-CoV-2 infection of HAEC culture.

**Fig. 2.**
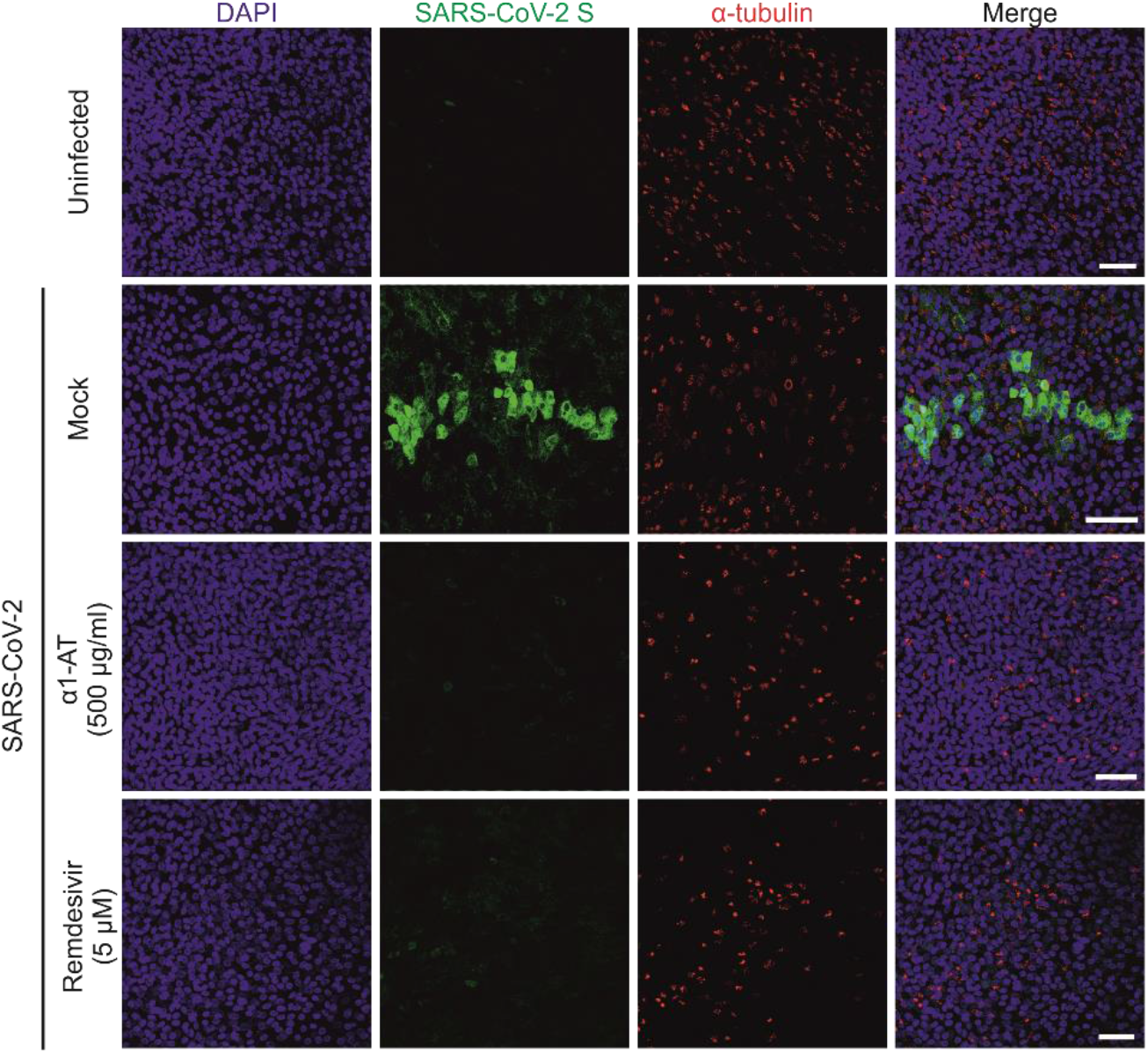
α1-AT inhibits SARS-CoV-2 infection of human airway epithelium cultures. The apical site of HAEC was exposed to buffer (mock), α1-AT (500 μg/ml) and remdesivir (5 μM) and then inoculated with SARS-CoV-2. Tissues were fixed 2 days later, stained with DAPI (cell nuclei), a SARS-CoV-2 specific spike antibody and an α-tubulin-specific antibody. Images represent maximum projections of serial sections along the basolateral to apical cell axis. Scale bar: 50 μm.

Our results demonstrate that endogenous lung- and plasma-derived α1-AT inhibit SARS-CoV-2 infection in cell lines and fully differentiated airway epithelium cultures. A recent preprint publication suggests that α1-AT may suppress the spike protein priming protease TMPRSS2, similar to Camostat mesylate^16^, however, evidence that α1-AT indeed inhibits SARS-CoV-2 infection was lacking. Our results using pseudoparticles and replication-competent virus are in line with this observation, because α1-AT prevents an early step in the viral life cycle, and selectively inhibits SARS-CoV-2 spike but not VSV-G mediated infection, which is independent from TMPRSS2 activation^17^. Thus, similar to Camostate mesylate, α1-AT may inhibit TMPRSS2-mediated priming of the Spike protein, thereby preventing engagement of the ACE2 receptor and subsequent fusion.

α1-AT has a reference range in blood of 0.9–2.3 mg/mL, and levels further increase 4-fold during acute inflammation. We show that plasma-derived α1-AT blocks SARS-CoV-2 infection with IC_50_ values ranging from ~ 0.5 to 1 mg/mL and almost completely inhibit virus infection at of 2-4 mg/mL. These concentrations are well within the range in blood and interstitial fluid^18^. Thus, α1-AT may serve as natural inhibitor of SARS-CoV-2 infection, in particular during acute infection when α1-AT concentrations increase. Thus, it will be interesting to further investigate whether the levels of α1-AT in the lungs or blood of COVID-19 patients inversely correlate with viral loads or disease progression.

Most importantly, α1-AT is an approved drug allowing its repositioning for the therapy of COVID-19. α1-AT deficiency is a hereditary disorder which results in reduced concentrations of the serpin and consequently a chronic uninhibited breakdown of tissue in the lungs. Several products containing α1-AT concentrated from human plasma (such as Prolastin® used herein) are approved since decades for intravenous augmentation therapy in patients suffering from α1-AT deficiency. Of note, α1-AT can also be administered at substantially higher doses than those in routine α1-AT deficiency (60 mg/kg/wk). Studies have administered α1-AT at 250 mg/kg without causing side effects and resulting in a 5-fold increase of the serpin concentrations in lung epithelial lining fluid^19^. Thus, rapid evaluation of α1-AT as drug in severe COVID-19 disease is highly warranted.

## Methods

### Ethic statements

Ethical approval for the generation of peptide libraries from lungs and BAL was obtained from the Ethics Committee of Ulm University (application numbers 274/12 and 324/12). The collection of tissue and generation of ALI cultures for research from these primary cells has been approved by the ethics committee at the University of Ulm (nasal brushings, application number 126/19) and Medical School Hannover (airway tissue, application number 2699-2015)

### Reagents

Camostat mesylate was obtained from Merck. Α1-antitrypsin was obtained from Merck or GRIFOLS (Prolastin®). EK1 was synthesized by solid phase synthesis.

### Generation of a peptide/protein library from lungs

6.5 kg of human lung were obtained from dead individuals without known diseases from pathology of Ulm University. The organs were frozen at −20°C and afterwards freeze dried. In the next step the lung material was homogenized and peptides and small proteins extracted by an ice-cold acetic acid extraction procedure. Then the mixture was centrifuged at 4.200 rpm, and the supernatant filtrated through a 0.45 μm filter. Thereafter, the obtained peptides and proteins were separated by ultrafiltration (cutoff:30kDa). The filtrate was then separated by reversed-phase (RP) chromatography with a Sepax Poly RP300 50×300mm (Sepax Technologies, Newark DE, USA 260300-30025) with a flow rate of 100 ml/min with the gradient program (min/%B): 0/5 5/5 20/25 35/50 50/100 being A, 0.1% TFA (Merck, 1082621000) in ultrapure water, and B, 0.1% TFA in acetonitrile (J.T.Baker, JT9012-3). Reversed-phase chromatographic fractions of 50 ml were collected to constitute the lung peptide bank, from which 1 mL-aliquots (2 %) were lyophilized and used for antiviral activity testing.

### Generation of a peptide/protein library from BAL

Clinical samples of BAL comprising a total of 20 L were collected and immediately frozen for further processing. Peptide/protein extraction was done by acidification with acetic acid to pH 3, followed by centrifugation at 4.200 rpm and filtration (0.45 μm) of the supernatant. Further, the filtered BAL was subjected to ultrafiltration (cut off: 30 kDa) yielding 22 L of a sample enriched in peptides and proteins. Chromatographic fractionation of the ultrafiltrated sample was performed by using a reversed-phase (PS/DVB) HPLC column Sepax Poly RP300 (Sepax Technologies, Newark DE, USA 260300-30025) of dimensions 3 x 25 cm, 121 at a flow rate of 55 mL/min with the gradient program (min/%B): 0/5 5/5 20/25 35/50 50/75 55/0, being A, 0.1% TFA (Merck, 1082621000) in ultrapure water, and B, 0.1% TFA in acetonitrile (J.T.Baker, JT9012-3). Seventy-three reversed-phase chromatographic fractions of 50 ml were collected to constitute the BAL peptide bank, from which 1 mL-aliquots (2 %) were lyophilized and used for antiviral activity testing. For further purification of active fractions, a second reversed-phase C18 HPLC 4,6×250mm column (Phenomenex 00G-4605-EO) at a flow rate of 0,8mL/min with gradient program (min/%B):0/5 50/60 60/100 was used.

### Cell culture

Unless stated otherwise, HEK293T cells were cultivated in DMEM supplemented with 10 % fetal calf serum (FCS), 2 mM L-glutamine, 100 U/ml penicillin and 100 mg/ml streptomycin. Caco2 cells were cultivated in DMEM supplemented with 10 % fetal calve serum (FCS), 2 mM Glutamine, 100 U/ml Penicillin and 100 mg/μl Streptomycin, 1x non-essential amino acids (NEAA) and 1 mM sodium pyruvate. TMPRSS2-expressing Vero E6 cells (kindly provided by the National Institute for Biological Standards and Control (NIBSC), #100978) were cultivated in DMEM supplemented with 10 % fetal calf serum (FCS), 2 mM L-glutamine, 100 U/ml penicillin, 100 mg/ml streptomycin and 1 mg/ml geneticin.

### Generation of lentiviral pseudotypes

For generation of lentiviral SARS-CoV-2 pseudoparticles (LV(Luc)-CoV-2-S) 900,000 HEK293T cells were seeded in 2 ml HEK293T medium. The next day, medium was replaced and cells were transfected with a total of 1 μg DNA using polyethylenimin (PEI). 2 % of pCG1-SARS-2-S (encoding the spike protein of SARS-CoV2 isolate Wuhan-Hu-1, NCBI reference Sequence YP_009724390.1, kindly provided by Stefan Pöhlmann) were mixed with pCMVdR8_91 (encoding a replication deficient lentivirus) and pSEW-Luc2 (encoding a luciferase reporter gene, both kindly provided by Christian Buchholz) in a 1:1 ratio in OptiMEM. Plasmid DNA was mixed with PEI at a DNA:PEI ratio of 1:3 (3 μg PEI per μg DNA), incubated for 20 min at RT and added to cells dropwise. At 8 h post transfection, medium was removed, cells were washed with 2 ml of PBS and 2 ml of HEK293T medium with 2.5 % FCS were added. At 48 h post transfection, pseudoparticles containing supernatants were harvested and clarified by centrifugation for 5 min at 1500 rpm.

### Generation of VSV-based pseudotypes

For generation of a VSV-based SARS-CoV-2 pseudoparticle (VSV(Luc_eGFP)-CoV-2-S), HEK293T were seeded in 30 ml HEK293T medium in a T175 cell culture flask. The next day, cells were transfected with a total 44 μg pCG1-SARS-2-S (encoding the pike protein of SARS-CoV2 isolate Wuhan-Hu-1, NCBI reference Sequence YP_009724390.1, kindly provided by Stefan Pöhlmann) using PEI. Plasmid DNA and PEI were mixed in 4.5 ml of OptiMEM at a 2:1 ratio (2 μg PEI per μg DNA), incubated for 20 min at RT and added to cells dropwise. 24 h post transfection, medium was replaced and cell were transduced with VSV-G-protein pseudotyped VSV encoding luciferase and GFP reporter gene (kindly provided by Gert Zimmer, Institute of Virology and Immunology, Mittelhäusern/Switzerland)^20^. At 2 h post transduction, cells were washed three times with PBS and cultivated for 16 h in HEPES-buffered HEK293T medium. Virus containing supernatants were then harvested and clarified by centrifugation for 5 min at 1500 rpm, residual pseudoparticles harboring VSV-G-protein were blocked by addition of anti-VSV-G hybridoma supernatant at 1/10 volume ratio (I1, mouse hybridoma supernatant from CRL-2700; ATCC). Virus stocks were concentrated 10-fold using a 100 kDa Amicon molecular weight cutoff and stored at −80°C until use.

### Screening lung and BAL library for inhibitors of SARS-CoV-2 infection

10,000 Caco2 cells (colorectal carcinoma cells) were seeded in 100 μl respective medium in a 96-well flat bottom plate. The next day, medium was replaced by 40 μl of serum-free medium. For screening peptide containing fractions, 10 μl of the solubilized fraction were added to cells. Cells were inoculated with 50 μl of infectivity normalized LV(Luc)-CoV2 (or LV(Luc)-no GP control). Transduction rates were assessed by measuring luciferase activity in cell lysates at 48 hours post transduction with a commercially available kit (Promega). Values for untreated controls were set to 100% infection.

### Gel electrophoresis and western blotting

Gel electrophoresis of active fractions was performed on a 4-12 % Bis-Tris protein gel (NuPAGE™) according to the manufacturers protocol. Prior to electrophoresis, samples were reduced by addition of 50 mM TCEP and heated for 10 min at 70 °C. Protein gel was either stained with Coomassie G-250 (GelCode™ Blue Stain) or applied to western blotting of α1-AT with a polyclonal anti-α1-AT antibody (Proteintech 16382-1-AP). The primary antibody was detected with labeled anti-rabbit secondary antibody and imaged with Odysey Infrared Imager (Licor).

### Tryptic in-gel digestion of proteins

Bands of interest were excised and the proteins were digested with trypsin. Tryptic peptides were eluted from the gel slices with 1% trifluoric acid.

### Matrix-assisted laser-desorption ionization time-of-flight mass spectrometry (MALDI-TOF-MS)

MALDI-TOF-MS was performed on an Ultraflex TOF/TOF mass spectrometer (Bruker Daltonics, Bremen) equipped with a nitrogen laser and a LIFT-MS/MS facility. The instrument was operated in the positive-ion reflectron mode using 2.5-dihydroxybenzoic acid and methylendiphosphonic acid as matrix. Sum spectra consisting of 200–400 single spectra were acquired. For data processing and instrument control the Compass 1.4 software package consisting of FlexControl 4.4, FlexAnalysis 3.4 4, Sequence Editor and BioTools 3.2 and ProteinScape 3.1. were used. External calibration was performed with a peptide standard (Bruker Daltonics).

### Database search

Proteins were identified by MASCOT peptide mass fingerprint search (http://www.matrixscience.com) using the Uniprot Human database (version 20200226, 210438 sequence entries; p<0.05). For the search, a mass tolerance of 75 ppm was allowed and oxidation of methionine as variable modification was used.

### Time-of addition experiments

One day prior to transduction, 10,000 Caco2 cells were seeded with respective medium in a 96 well plate. For the addition of α1-AT prior to infection, medium was replaced by 80 μl of serum-free medium and cells were incubated with serial dilutions of α1-AT for 0, 1, 2 and 4 h at 37 °C followed by infection with 20 μl of infectivity normalized VSV(Luc)-CoV-2-S pseudoparticles. To investigate whether α1-AT acts post viral entry, medium was replaced by 100 μl of serum-free medium. Cells were inoculated with 20 μl of infectivity normalized VSV(Luc)-CoV-2 pseudoparticles and incubated for 2 h at 37 °C. After incubation period cells were washed with 100 μl PBS and 100 μl serum-free medium as well as 20 μl of serially diluted α1-AT were added. Infection rates were assessed by measuring luciferase activity in cell lysates at 16 h post transduction with a commercially available kit (Promega). Values for untreated controls were set to 100% infection.

### Cell viability assay

To assess cytotoxicity of α1-AT and camostat mesylate, 10,000 Caco2 cells were seeded in 100 μl medium in a 96-well flat bottom plate. The next day, medium was replaced by 80 μl of serum-free Caco2 medium and cells were treated with serial dilutions of Prolastin®, camostat mesylate or DMSO as solvent control for camostat mesylate. After 48 h, cell viability was assessed by measruring ATP levels in cells lysates with a commercially available kit (CellTiter-Glo®, Promega).

### MTS cell viability assay

To quantify SARS-CoV-2 wildtype infection, virus induced cell death was inferred from remaining cell viability determined by MTS (3-(4,5-dimethylthiazol-2-yl)-5-(3-carboxymethoxyphenyl)-2-(4-sulfophenyl)-2H-tetrazolium) assay. To this end, 20,000 TMPRSS2-expressing Vero E6 cells were seeded in 96 well plates in 100 μl respective medium. The next day, medium was replaced with 132 μl serum-free medium and the respective compound of interest was added. After incubation for 1 h at 37°C the cells were infected with a multiplicity of infection of 0.001 (MOI; based on PFU per cell) of the viral isolates BetaCoV/France/IDF0372/2020 (#014V-03890) and BetaCoV/Netherlands/01/NL/2020 (#010V-03903), which were obtained through the European Virus Archive global, in a total volume of 180 μl. 2 days later, infection was quantified by detecting remaining metabolic activity. To this end, 36 μl of CellTiter 96® AQueous One Solution Reagent (Promega G3580) was added to the medium and incubated for 3 h at 37°C. Then, optical density (OD) was recorded at 620 nm using a Asys Expert 96 UV microplate reader (Biochrom). To determine infection rates, sample values were subtracted from untreated control and untreated control set to 100%.

### Generation of human airway epithelial cells

Differentiated ALI cultures of human airway epithelial cells (HAECs) were generated from primary human basal cells isolated from airway epithelia. Cells were expanded in a T75 flask (Sarstedt) in Airway Epithelial Cell Basal Medium supplemented with Airway Epithelial Cell Growth Medium SupplementPack (both Promocell). Growth medium was replaced every two days. Upon reaching 90 % confluence, HAECs were detached using DetachKIT (Promocell) and seeded into 6.5 mm Transwell filters (Corning Costar). Filters were precoated with Collagen Solution (StemCell Technologies) overnight and irradiated with UV light for 30 min before cell seeding for collagen crosslinking and sterilization. 3.5 x 10^4^ cells in 200 μl growth medium were added to the apical side of each filter, and an additional 600 μl of growth medium was added basolaterally. The apical medium was replaced after 48 h. After 72 – 96 h, when cells reached confluence, the apical medium was removed and basolateral medium was switched to differentiation medium ± 10 ng/ml IL-13 (IL012; Merck Millipore). Differentiation medium consisted of a 50:50 mixture of DMEM-H and LHC Basal (Thermo Fisher) supplemented with Airway Epithelial Cell Growth Medium SupplementPack and was replaced every 2 days. Air-lifting (removal of apical medium) defined day 0 of air-liquid interface (ALI) culture, and cells were grown at ALI conditions until experiments were performed at day 25 to 28. To avoid mucus accumulation on the apical side, HAEC cultures were washed apically with PBS for 30 min every three days from day 14 onwards.

### Effect of remdesivir and α1-antitrypsin on SARS-CoV-2 infection of HAECs

Immediately before infection, the apical surface of HAECs grown on Transwell filters were washed three times with 200 μl PBS to remove accumulated mucus. Then, 5 μM remdesivir were added into the basal medium or 500 μg/ml α1-AT were added into the basal medium and onto the apical surface. Cells were infected with 9.25 × 10^2^ plaque forming units (PFU) of SARS-CoV-2 (BetaCoV/France/IDF0372/2020). After incubation for 2 h at 37°C, viral inoculum was removed and cells were washed three times with 200 μl PBS. At 2 days post infection, cells were fixed for 30 min in 4 % paraformaldehyde in PBS, permeabilized for 10 min with 0.2% saponin and 10% FCS in PBS, washed twice with PBS and stained with anti-SARS-CoV-2 (1A9; Biozol GTX-GTX632604) anti-MUC5AC (clone 45M1; MA1-21907, Thermo Scientific) and anti-alpha-tubulin (MA1-8007, Thermo Scientific diluted 1:300 to 1:500, respectively, in PBS, 0.2% saponin and 10% FBS over night at 4 °C. Subsequently cells were washed twice with PBS and incubated for 1 h at room temperature in PBS, 0.2% saponin and 10% FCS containing AlexaFluor 488-labelled anti-rabbit, AlexaFluor 468-labelled anti-mouse and AlexaFluor 647-labelled anti-rat secondary antibody, respectively (all 1:500; Thermo Scientific) and DAPI + phalloidin AF 405 (1: 5,000; Thermo Scientific). Images were taken on an inverted confocal microscope (Leica TCS SP5) using a 40x lens (Leica HC PL APO CS2 40×1.25 OIL). Images for the blue (DAPI), green (AlexaFluor 488), red (AlexaFluor 568) and far-red (AlexaFluor 647) channels were taken in sequential mode using appropriate excitation and emission settings.

### Non-linear regression and statistics

Analysis was performed using GraphPad Prism version 8.4.2. Calculation of IC_50_ values via non-linear regression was performed using normalized response-variable slope equation. For statistical analysis, a 2-way ANOVA with Dunett’s multiple comparison test was used.

## Data availability

All primary data will be made accessible to qualified researchers upon request.

## Acknowledgments

L.W., C.C., T.W., R.G., M.H., A.S. are part of and R.G. is funded by a scholarship from the International Graduate School in Molecular Medicine Ulm. This work was supported by the EU’s Horizon 2020 research and innovation programme (Fight-nCoV, 101003555 to J.M.) and by the DFG (CRC1279 to S.S., F.K. and J.M.).

## Author contributions

L.W. generated LV pseudotypes, performed screening, gel electrophoresis, and a1-AT inhibitions studies with pseudotypes; C.C. and J.A.M generated SARS-CoV-2 stocks und performed all infection experiments in the BSL-3; F.Z., C.P.B, R.G. and A.S. generated VSV pseudotypes; T.W. supported L.W. in most experiments; M.H. performed Western Blots; S.K supported L.W. in screening; N.P. synthesized EK1, generated peptide libraries and purification; G.F. and R.B. generated HAEC and did stainings; P.K. and V.M. provided reagents and controls; L.S. supervised the generation of peptide libraries; D.R.T. performed autopsies and collected lungs; C.S. collected BAL; S.S. supervised BSL-3 work; A.K. advised and wrote the manuscript; G.L performed mass spectrometry; F.K. supervised work and wrote the manuscript; M.F. supervised work with HAEC; J.M. is responsible for the study, supervised all work and wrote the manuscript;

## Competing interests

The authors declare no competing interests.

## Additional information

**Extended data Fig. 1.**
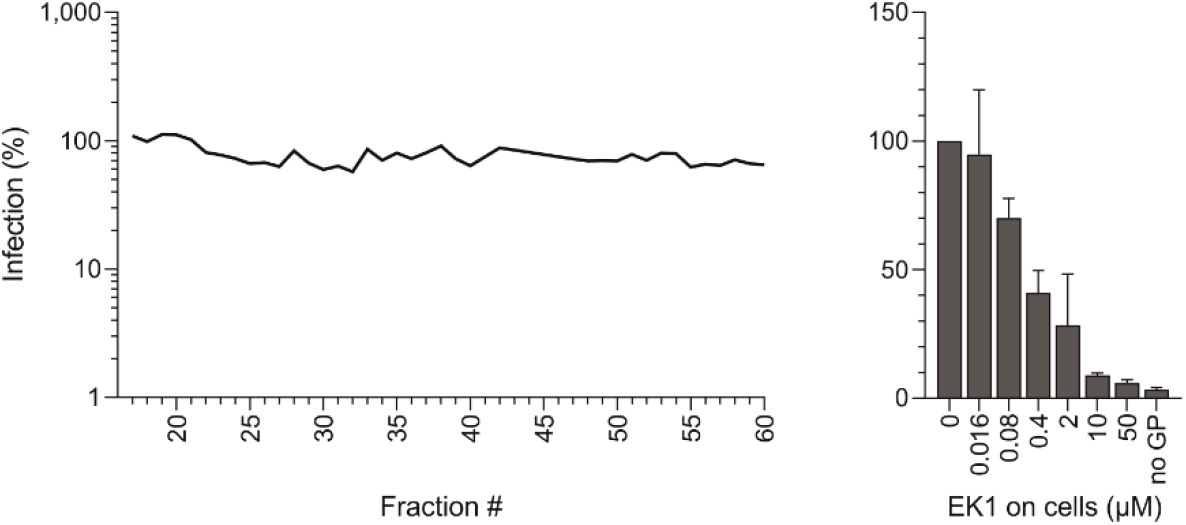
Screening a peptide/protein library made from lungs for fractions with activity against SARS-CoV-2. The 39 fractions were added to Caco2 cells and infected with SARS-CoV-2 Spike pseudoparticles. Infection was determined two days later by luciferase assay. The spike specific peptide entry blocker EK1 was used as control. Data shown were derived from duplicate (screen) and triplicate infections (EK1).

**Extended data Fig. 2.**
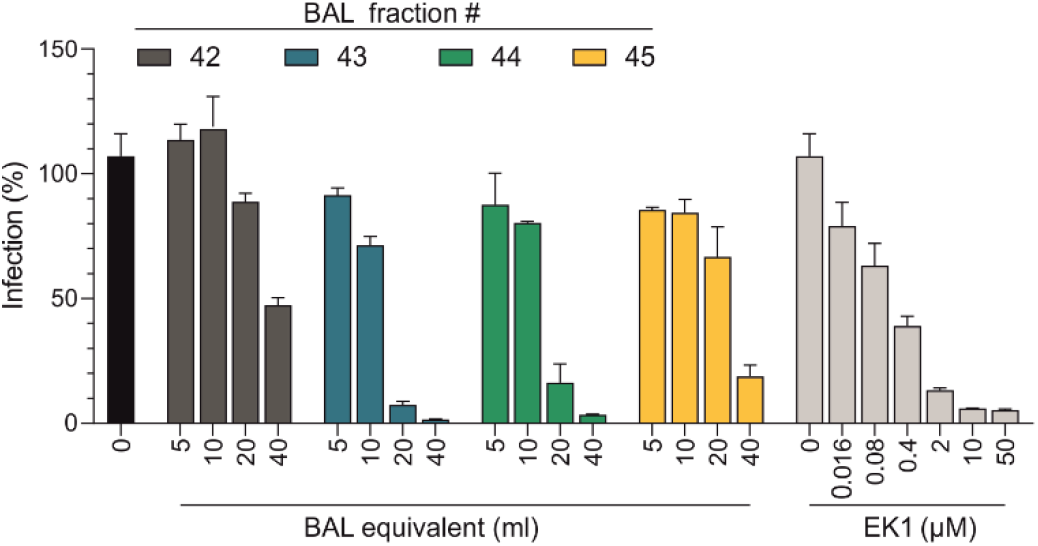
BAL fractions 42-45 inhibit SARS-CoV-2 pseudotype infection. Serial dilutions of the BAL fractions 42 to 45 and EK1 (see Fig. 1a) were added to Caco2 cells, which were subsequently infected with SARS-CoV-2 spike pseudoparticles. Infection rates were determined two days later by luciferase assay. Data shown were derived from and triplicate infections.

**Extended data Fig. 3.**
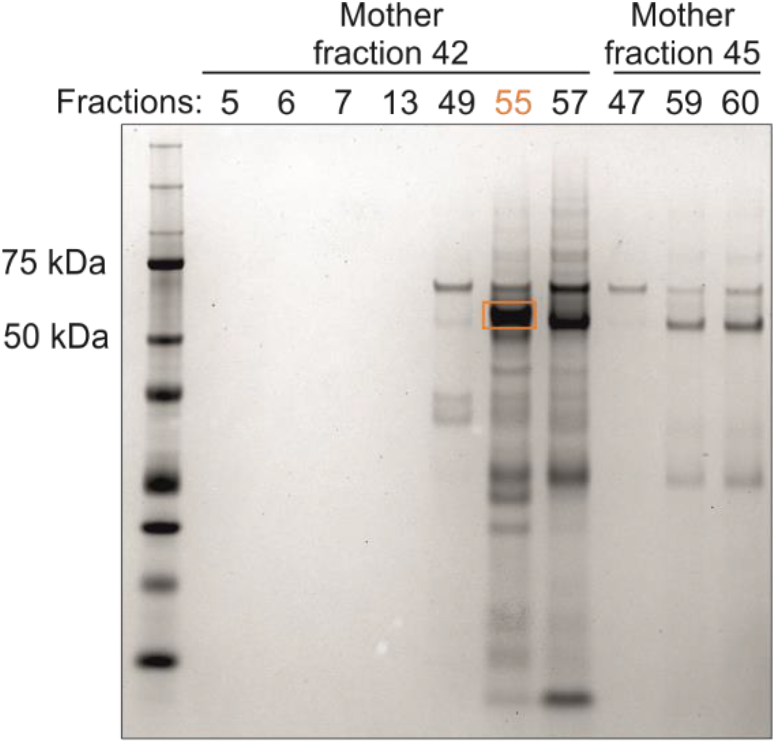
Gel electrophoresis of BAL fractions. Gel electrophoresis of active fractions was performed on a 4-12 % Bis-Tris protein gel. Prior to electrophoresis, samples were reduced by addition of 50 mM TCEP and heated for 10 min at 70 °C. Protein gel was stained with Coomassie G-250. The boxed band fraction 42/55 was cut out and subjected in to an in gel tryptic digest.

**Extended data Fig. 4.**
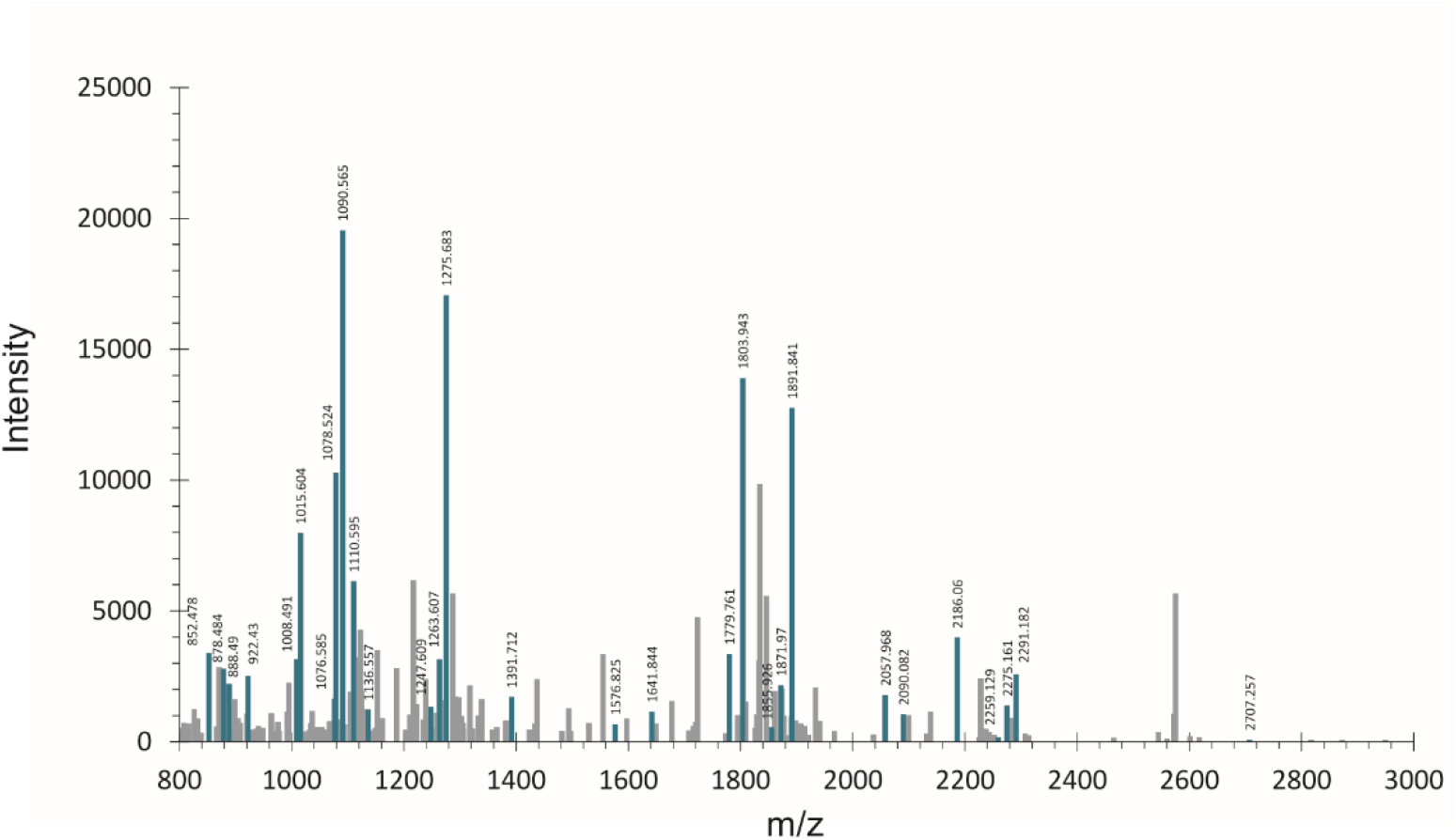
**MALDI-TOF-MS spectrum** of the tryptic digest of boxed band from gel shown in Extended data Fig. 3. Mass signals assigned to the identified alpha-1 antitrypsin are shown in green. Also see extended table 1.

**Extended data Fig. 5.**
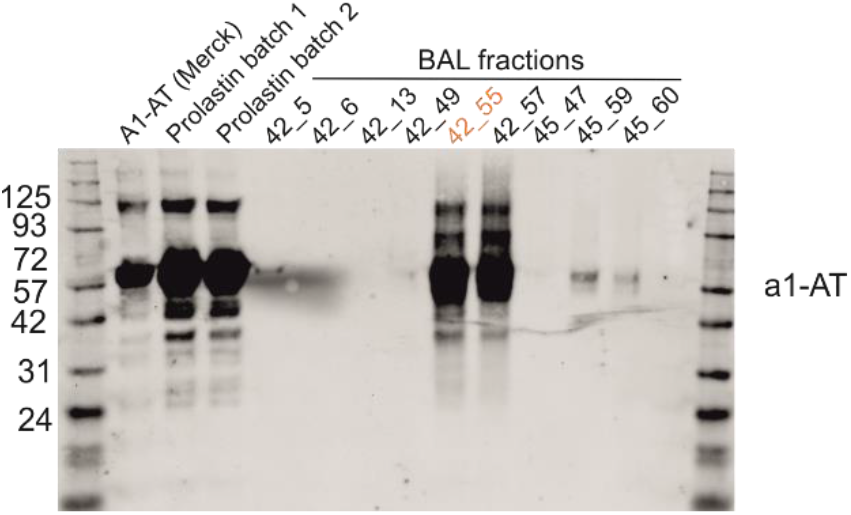
Western Blot analysis of α1-antitrypsin samples and BAL fractions.

**Extended data Fig. 6.**
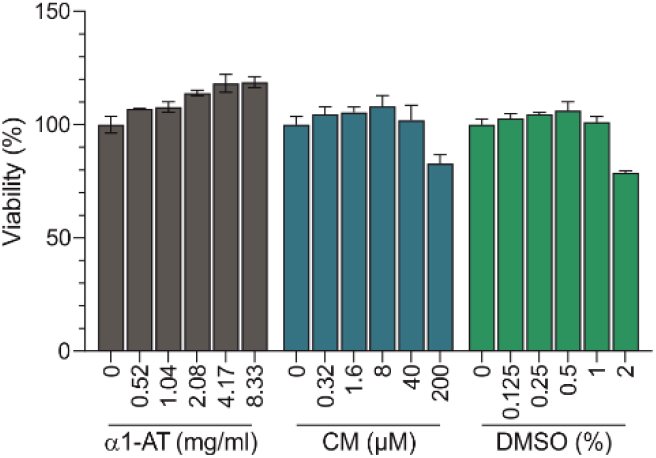
Cell viability assay. To assess cytotoxicity of α1-AT and Camostat mesylate (CM), Caco2 cells were treated with serial dilutions of the compounds (and DMSO as solvent control for CM). After 48 h, cell viability was assessed by measuring ATP levels in cells lysates with a commercially available kit (CellTiter-Glo®, Promega).

**Extended data Fig. 7.**
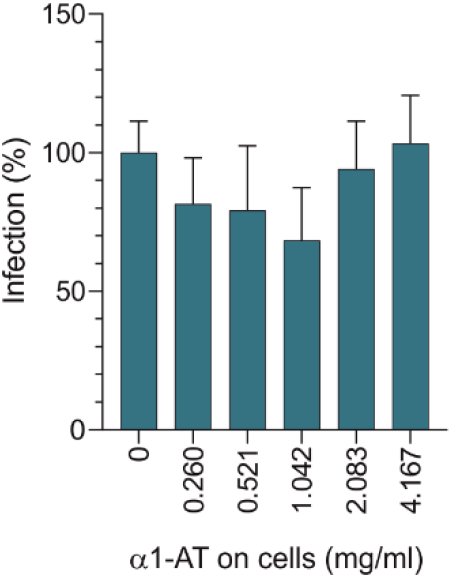
α1-AT has no effect on SARS-CoV-2 spike pseudoparticle infection if added post entry. Caco2 cells were infected with SARS-CoV-2 spike pseudoparticles. After 2 hours, inoculum was removed and α1-AT was added. Infection rates were determined 2 dpi by measuring luciferase activities in cellular lysates. Values shown were derived from triplicate infections and show mean values ± sd.

**Extended data Fig. 8.**
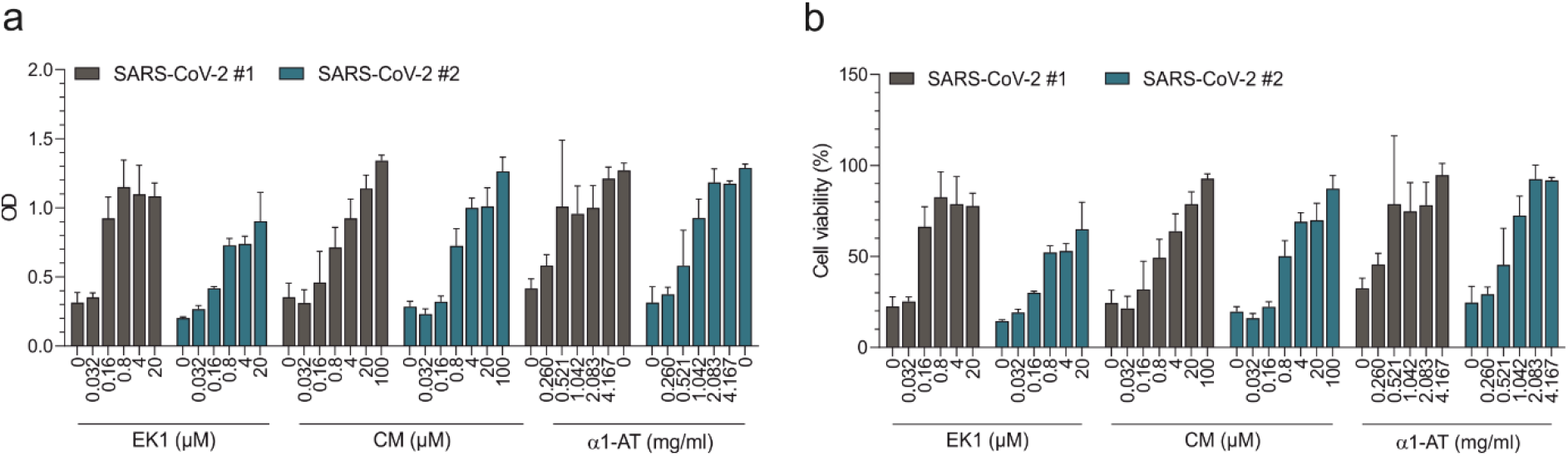
α1-AT inhibits SARS-CoV-2 infection. TMPRSS2-expressing Vero cells were exposed to indicated concentrations of EK1, CM and α1-AT and then infected with a French (#1) and Dutch (#2) SARS-CoV-2 isolate. 2 days later, virus induced cytopathic effects were determines by XTS assay. a) Optical density (OD) was recorded at 620 nm using a Asys Expert 96 UV microplate reader (Biochrom). b) To determine infection rates, sample values were subtracted from untreated control and untreated control set to 100%. Values shown were derived from triplicate infections and show mean values ± sd.

**Supplementary Table 1.**
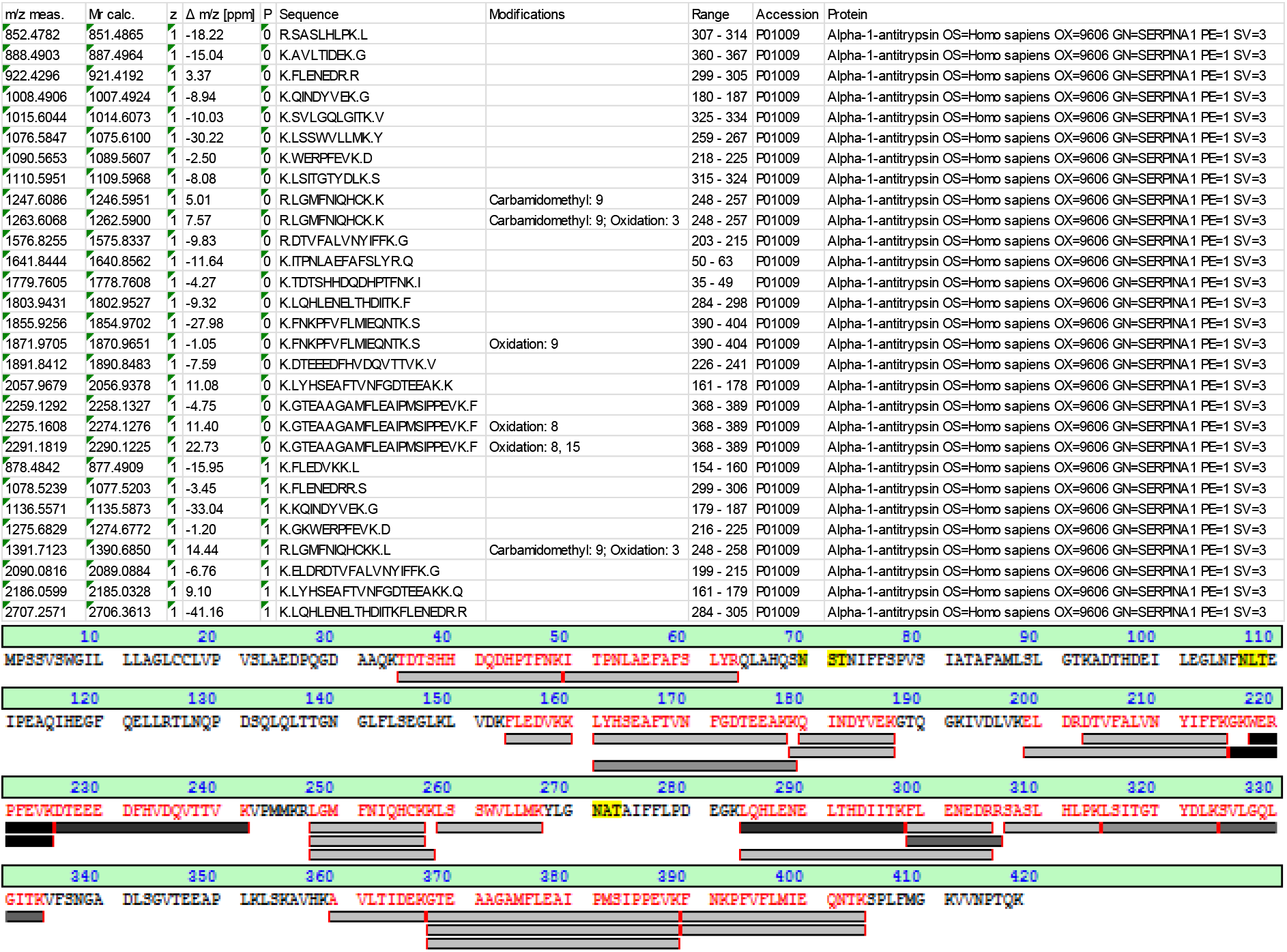

## Notes

### Competing Interest Statement

The authors have declared no competing interest.

